# Cell-type-specific encoding of effort-based decision-making in the dorsal raphe nucleus

**DOI:** 10.64898/2026.09.28.751674

**Authors:** Laura Chalençon, Robert N Fetcho, Alexandra Lenz, Joseph M Stujenske, Matthew A Wright, Conor Liston

## Abstract

Effort-based decision-making, the process of weighing reward benefits against effort costs, is essential for adaptive behavior. Although the dorsal raphe nucleus (DRN) contributes to reward processing and effort-related behaviors, how specific DRN neuronal subtypes contribute to effort-based decision-making remains poorly understood. Using fiber photometry to record DRN serotonergic and GABAergic neurons during an effort-based decision-making task, we found that tonic fluctuations in DRN serotonergic activity were selectively associated with persistent selection of the high-value option. Furthermore, we found that while neither neuronal population was sensitive to relative reward magnitude, effort expenditure selectively enhanced decision- and reward-related signals carried by serotonergic and GABAergic neurons, respectively. Together, these findings support a cell type-specific model in which serotonergic activity provides a signal linking persistent motivational states to effortful choice, whereas GABAergic activity may contribute to evaluating or updating reward outcomes after effort expenditure.

**SIGNIFICANCE STATEMENT:** Choosing whether a reward is worth the effort required to obtain it is essential for adaptive behavior, yet the neural mechanisms underlying this process remain incompletely understood. Here, we identify distinct contributions of serotonin- and GABAergic neurons in the dorsal raphe nucleus, a major source of brain serotonin, to effort-based decision-making. Serotonergic neurons were preferentially engaged in sustained motivational state and action selection, and GABAergic neurons in the processing of reward outcomes following costly actions. Our findings reveal a cell type-specific organization of effort-based reward seeking in the dorsal raphe nucleus and provide a framework for understanding how motivational states, decisions, and reward outcomes are coordinated.

## Introduction

Effort-based decision-making, the process of weighing reward benefits against effort costs^1,2^ is essential for adaptive behavior. Impairments in this process are associated with motivational deficits in neuropsychiatric disorderssuch as depression^2–4^, schizophrenia^5^, Parkinson’s disease^6,7^, and substance use disorders^8^. However, the neural mechanisms underlying effort-based decision-making remain incompletely understood.

The dorsal raphe nucleus (DRN) is a key regulator of adaptative behavior. The DRN, which constitutes the main source of forebrain serotonin, contains both serotonergic and GABAergic neurons. These two populations display marked molecular heterogeneity^9^ and differ in their anatomical organization and connectivity patterns^10^. While serotonergic neurons project extensively throughout the brain, providing widespread modulatory influences ^11^, GABAergic neurons contribute both to local microcircuit regulation within the DRN^12,13^ and to long-range projections targeting extra-raphe structures^14^. In addition, the two populations receive inputs from partially segregated neural circuits^10^, further supporting their engagement in different circuit functions. Consistent with their molecular heterogeneity and distinct circuit organization, evidence suggests that serotonergic and GABAergic neurons exert both complementary and opposing influences on behavior depending on the behavioral context^15–24^.

Theoretical models of effort-based decision-making posit the involvement of multiple brain regions and circuits encoding different types of effort signals and outcomes for regulating behavior to maximize reward^25,26^. Consistent with these frameworks, several brain regions, including the prefrontal cortex, striatum, hippocampus, and amygdala have been implicated in effort-based decision-making^26–32^. In contrast, despite its established role in behavioral regulation, the contribution of the DRN to effort-based decision-making remains incompletely understood.

Previous studies have established the DRN as a key component of reward-related processing. DRN neurons exhibit reward-related activity in both a projection-specific^33^ and cell-type-specific manner^34–36^. These reward-related signals can occur on distinct timescales, including rapid phasic responses to reward delivery^33^, as well as slower tonic signals that may encode more sustained aspects of reward or motivational state^34,37^. Beyond reward encoding, DRN activity and serotonergic signaling influence several dimensions of cost–benefit evaluation, including probability and risk^38–41^ assessment, and delay discounting^42^. In parallel, DRN circuits, including serotoninergic and GABAergic neurons, contribute to effort-related behavior in a cell-type specific manner^22,43^. These observations raise the possibility that the DRN may contribute to effort-based decision-making by encoding both reward value and effort cost.

Consistent with this hypothesis, pharmacological studies targeting serotonergic signaling have shown that administration of the 5-HT1A receptor agonist 8-OH-DPAT biases rats toward low-effort, low-reward choices^44^. In humans, enhancement of serotonergic transmission via the selective serotonin reuptake inhibitor escitalopram has been reported to increase the physical force healthy participants are willing to exert to obtain monetary rewards^45^. However, the mechanisms by which DRN neurons contribute to effort-based decision-making remain poorly understood, particularly with respect to their distinct cellular subtypes. In particular, it remains unclear whether the coding properties of different features of effortful reward seeking are subtype-specific or shared between DRN serotonergic and GABAergic neuronal populations.

## Results

To assess the involvement of the DRN in effortful reward-seeking, mice were trained in an effort-based decision-making task in which choices were guided by both reward magnitude and effort cost. As previously described^29,46,47^, freely moving mice learned to navigate a T-maze by turning left or right to obtain a low or a high reward (water of varying volumes). After learning this contingency, an individual’s willingness to expend effort for a larger reward was evaluated by introducing a physical barrier that must be climbed to obtain the high-reward option. Mice were trained through a series of session types based upon the amount of effort needed to acquire the high-reward option: no-effort, low-effort and high-effort. Importantly, while the low-reward option was never associated with an effort requirement, the high-reward option was successively paired with no-effort, low-effort or high-effort expenditures (no barrier, low barrier or high barrier) (**Figure 1A-B**). On each session, reward-seeking behavior was quantified as the percentage of high-reward choices during a session. The ratio between high- and low-reward volumes was individually adjusted to ensure that mice maintained a preference for the high-reward option. When the high-reward option was chosen on more than 70% of trials for two sessions in a row, animals progressed to the next type of session involving a new effort condition. Mice that were initially unwilling to exert effort for the larger reward progressively adapted their behavior as the reward ratio was adjusted. Consequently, animals displayed a stronger preference for the high-reward option during the final compared with the initial session of each effort condition (**Figure 1C**). Moreover, high-reward choices were not randomly distributed throughout sessions but instead occurred in temporally clustered sequences (**Figure 1D-E**). These clusters of consecutive choices toward the same reward option reflect a form of choice persistence, whereby animals repeatedly exploit a preferred option over successive trials.

**Figure 1:**
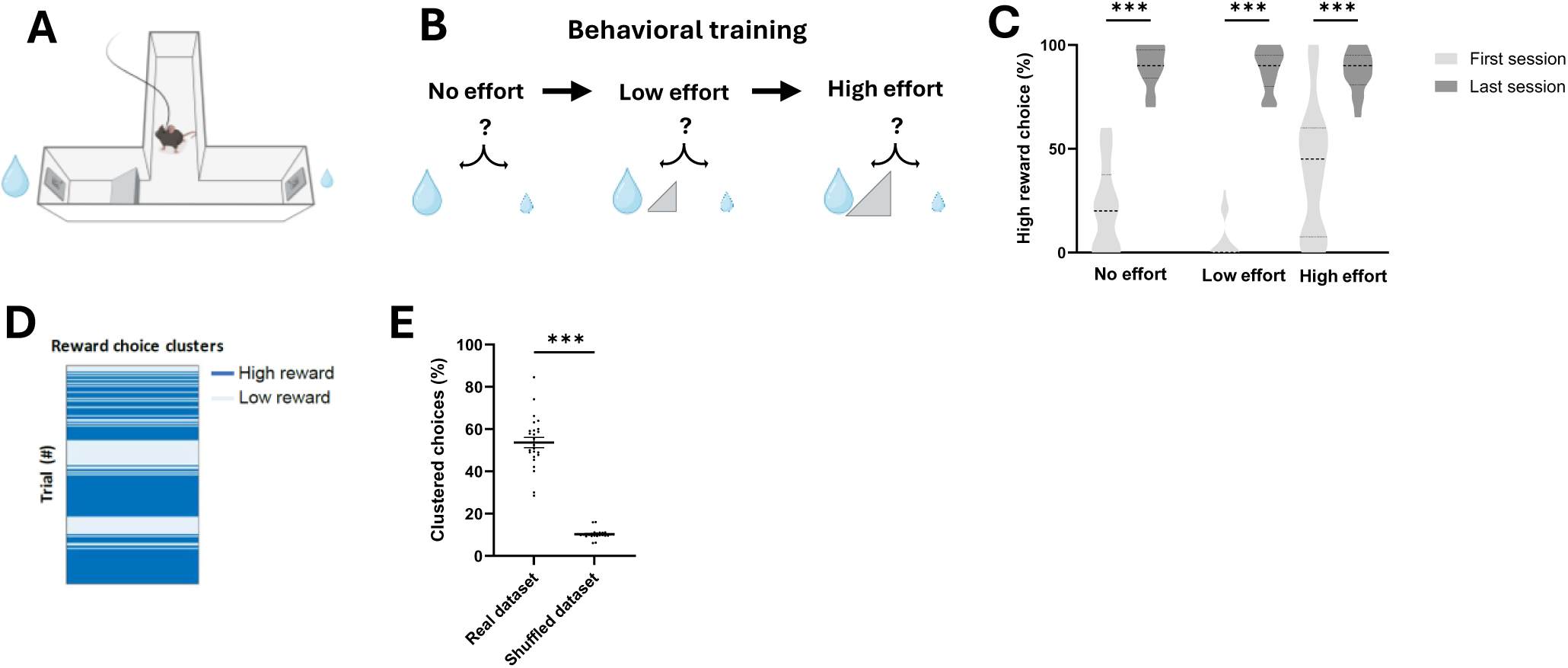
Effort-based decision-making task. **A.** Schematic of the effort-based decision-making behavioral apparatus. **B.** Timeline of the training protocol. Animals first learned to discriminate between the high- and low-reward options (no effort sessions). The high-reward option was then successively associated with increasing effort requirements (low effort, and high effort session). **C.** Mice preference for the high reward option during the first and the last session of no effort, low effort or high effort sessions (n=25 mice, Mixed-effects analysis: Session effect p<0.001, Effort effect p<0.001, Session*Effort effect p<0.001, Bonferroni corrected post-hoc for first vs last session: NE p<0.001, LE p<0.001, HE p<0.001). **D.** Example of the distribution of reward choices across all sessions for a representative mouse, illustrating the clustering of reward choices. **E.** The percentage of clustered trial is above chance level (n=25 mice, t=17.12 p<0.001). Violin plots represent mean (dashed lines), first and third quartiles (dotted lines).

To distinguish the contributionsof GABAergic vs. serotonergic neuronal populations of the DRN in the effort-based decision-making task, we used fiber photometry to specifically record the activity of serotonergic or GABAergic neuronal populations during task performance. To this end, we injected an inducible genetically encoded calcium indicator (GCaMP6s) into the DRN of SERT-flp or Gad2-cre mice respectively. We then implanted an optical fiber into the DRN to directly record the activity of either DRN^SERT^ or DRN^Gad2^ neurons (**Figure 2A-B**, **Supplementary figure 1 A-B**). The recording revealed slow fluctuations in tonic activity associated with rapid transient phasic responses to task events in both neuronal populations (**Figure 2C**). As tonic DRN activity has been implicated in reward-seeking behavior^34^, we first performed spectral analyses of DRN^SERT^ and DRN^Gad2^ activity to characterize slow fluctuations associated with tonic signaling during task performance. Spectral analyses revealed that activity in both populations was dominated by slow delta-band activity, with smaller contributions from theta- and alpha-band frequencies (**Figure 2D**, **Supplementary figure 1D-E**). Notably, DRN^SERT^ neurons exhibited stronger delta-band activity than DRN^Gad2^ neurons (**Figure 2E**).

**Figure 2:**
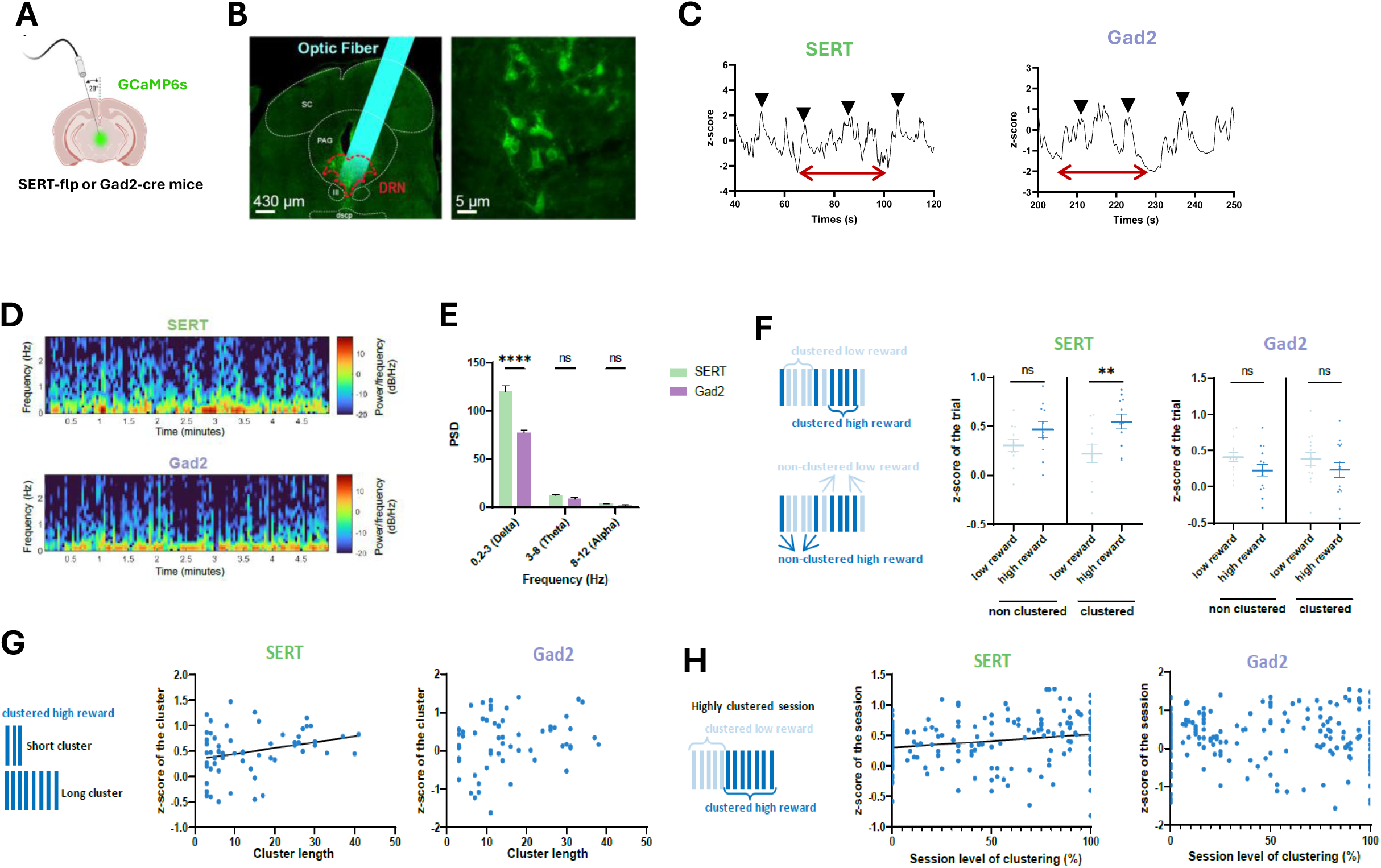
DRN^SERT^ and DRN^GAD2^ tonic activity during effort-based decision-making. **A.** Schematic of virus injection and fiber placement for fiber photometry recording of DRN neurons. **B.** Representative histological image of DRN showing GCaMP6s expressing cell bodies (green) in DRN with fiber-optic implant. DRN: Dorsal raphe nucleus (red dotted line). SC: Superior colliculus, PAG: Periaqueductal gray, III: Oculomotor nucleus, dscp: superior cerebellar peduncle decussation (white dotted line). **C.** Example traces of SERT (**left**) and Gad2 (**right**) recordings. Red arrows indicate slow tonic fluctuations, and black triangles indicate rapid phasic responses to task events. **D.** Spectrogram focusing on the low frequency bands of one example session of SERT (**top**) and Gad2 (**bottom**) recording. **E.** Gad2 neurons show less power spectral density (PSD) in the delta band than SERT neurons (n=11 SERT mice, n=14 Gad2 mice, Mixed-effects analysis: Group effect p<0.001, Frequency band effect p<0.001, Group*Frequency band effect p<0.001, Bonferroni corrected post-hoc for SERT versus Gad2: Delta band p<0.001, Theta band p>0.999, Alpha band p>0.999). **F.** SERT (**left**) and Gad2 (**right**) mean activity of high or low reward trials. SERT activity is higher during high reward trials in comparison to low reward trials, only when reward choices are clustered (n=11 mice, Mixed-effects analysis: Cluster effect p=0.973, Reward effect p=0.008, Cluster*Reward effect p=0.087, Bonferroni corrected post-hoc for high versus low reward trials: Clustered p=0.002, Non clustered p=0.148). There is no difference in the mean trial activity of Gad2 neurons between high or low reward trials, whether choices are clustered or not (n=14 mice, Mixed-effects analysis: Cluster effect p=0.906, Reward effect p=0.062, Cluster*Reward effect p=0.795, Bonferroni corrected post-hoc for high versus low reward trials: Clustered p=0.243, Non clustered p=0.143). **G.** Correlation between SERT (**left**) and Gad2 (**right**) mean activity of cluster of high reward choices and the number of trials constituting the cluster. SERT activity of a cluster of high reward choices is positively correlated with the length of the cluster (n=61 clusters, Simple linear regression, Pearson’s r=0.28 p=0.029). There is no correlation in Gad2 neurons (n=61 clusters, Simple linear regression, p=0.699, one outlier point has not been plotted). **H.** Correlation between SERT (**left**) and Gad2 (**right**) mean activity of the session and the percentage of clustered high reward choices over the total number of high reward choices of the session. Mean session activity of SERT neurons is positively correlated with the level of clustering of the session (n=194 sessions, Simple linear regression, Pearson’s r=0.20 p=0.007). No correlation is found in Gad2 neurons (n=224 sessions, Simple linear regression, p=0.982). Bar plots represent mean ±SEM. Swarm plots represent mean (line) ±SEM and individual datapoints (dots). Scatter plots represent individual datapoints (dots).

Because these slow fluctuations are thought to reflect tonic variations in population activity^48^, we next examined whether they were related to behavioral choice patterns using three complementary approaches. First, we asked whether tonic neuronal activity varied with high-reward choice persistence by comparing activity during trials in which mice chose to consume the high reward versus the low reward as a function of whether those choices were clustered or isolated during a given session (**Figure 2F**). Clustered trials were defined as sequences of consecutive choices toward the same reward option that were longer than expected by chance, based on a mouse-specific threshold derived from shuffled data. In DRN^SERT^ neurons, mean trial activity was higher during clusters of high-reward choices than during clusters of low-reward choices. This difference was not significant when the choices were not clustered, suggesting a cumulative effect of choice persistence on DRN^SERT^ neuronal activity. In contrast, no change in GABA signaling was observed between high reward and low reward choices, whether the choices were clustered or not. Furthermore, DRN^SERT^ activity during high-reward choice clusters positively correlated with cluster length, further suggesting that DRN^SERT^ activity scales with the degree of behavioral persistence (**Figure 2G**). This correlation was not observed in DRN^Gad2^ population. Finally, we neuronal activity across a session varied with behavioral choice persistence by analyzing the mean activity of the session as a function of the level of clustering of high-reward choices in that session. We found a positive correlation between the SERT signal and the level of clustering in a given session, but no association for the GABA signal (**Figure 2H**). Together, these findings indicate that both DRN^SERT^ and DRN^Gad2^ populations exhibit slow fluctuation of activity during effortful reward seeking, but only tonic fluctuations in DRN^SERT^ activity are associated with decision making-related behavior.

To further characterize the coding properties of effort-based decision-making by DRN^SERT^ and DRN^Gad2^ neurons, we examined whether these two neuronal populations exhibited phasic changes in activity in response to different behavioral components of the task, and whether these phasic responses were subtype-specific. We identified three behavioral epochs of interest that occur during the course of a trial—trial start, decision making (before turning left or right), and reward acquisition (**Figure 3A**)—and quantified SERT and GABA activity around these periods (**Figure 3B**). DRN^SERT^ and DRN^Gad2^ neurons displayed distinct patterns of activity across behavioral epochs (**Figure 3C**). Indeed, while the SERT signal decreased at the beginning of the trial (**Figure 3D**) and exhibited a clear increase time locked to the decision-making process (**Figure 3E**), the GABA signal increased at the beginning of the trial (**Figure 3D**) and did not vary during decision making process (**Figure 3E**). Both SERT and GABA signals exhibited a marked decrease time locked to the acquisition of the reward (**Figure 3F**). These phasic responses were consistent across trials (**Figure 3G-I**). These findings reveal cell type-specific phasic responses across distinct stages of the decision-making process.

**Figure 3:**
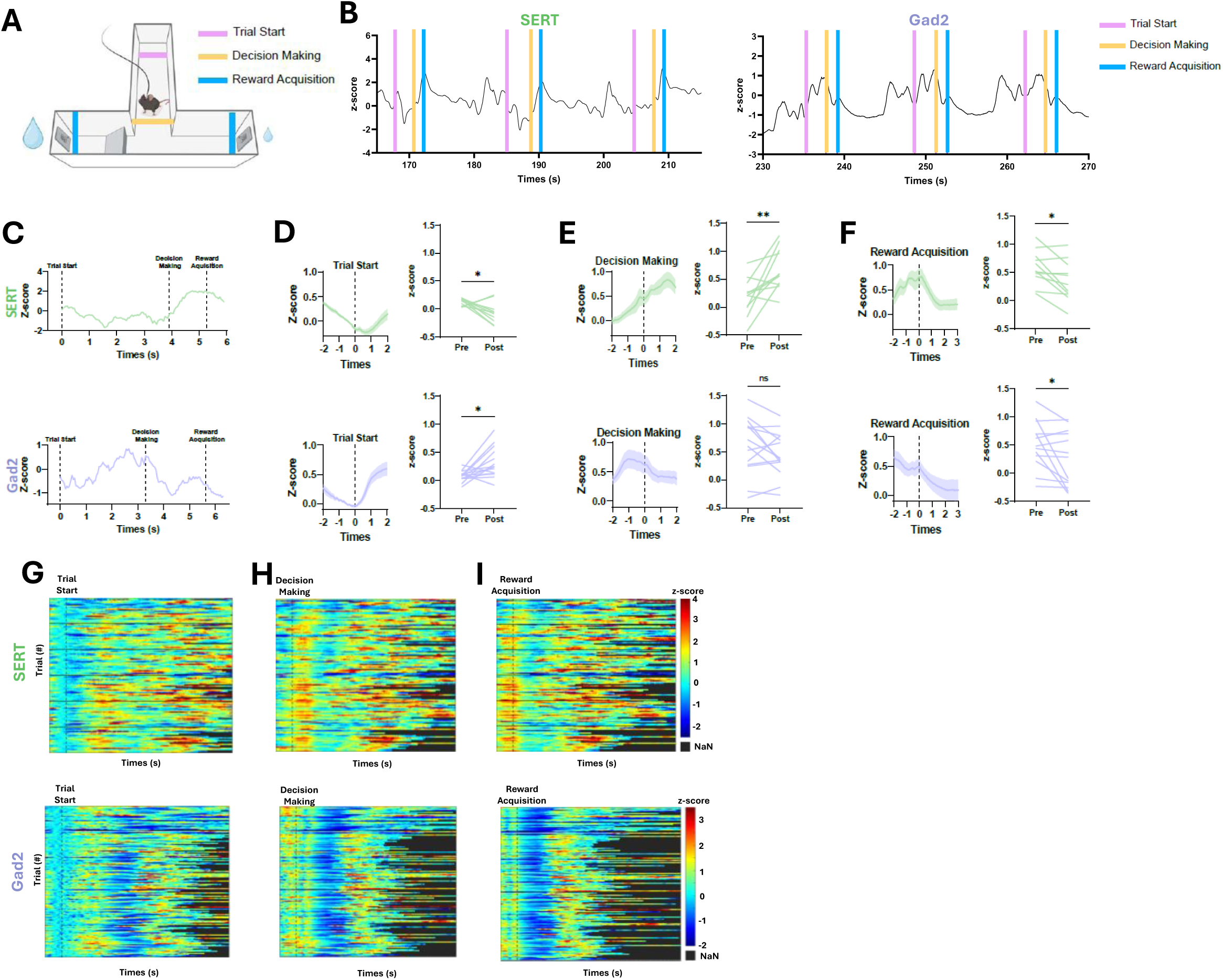
Phasic responses of DRN^SERT^ and DRN^Gad2^ neurons across task epochs. **A.** Schematic of the behavioral apparatus showing the task epochs used for analysis (Trial Start, Decision Making, and Reward Acquisition). **B.** Representative traces of SERT (**up**) and Gad2 (**down**) recoding across several trials. **C.** Representative traces of SERT (**up**) and Gad2 (**down**) during one no-effort trial. **D-F.** Averaged traces of SERT (**up**) and Gad (**down**) activity during no-effort sessions time-locked to Trial Start (**D**), Decision Making (**E**), and Reward Acquisition (**F**), and quantification of the signal changes pre to post each epoch. SERT activity is decreasing after Trial Start, increasing after Decision-Making and decreasing after Reward Acquisition (n=11 mice, Wilcoxon tests: Trial Start W=-52 p=0.019, Decision Making W=56 p=0.010, Reward Acquisition W=-52 p=0.019). Gad2 activity is increasing after Trial Start, doesn’t change after Decision-Making and is decreasing after Reward Acquisition (n=14 mice, Wilcoxon tests: Trial Start W=79 p=0.011, Decision Making W=-43 p=0.194, Reward Acquisition W=-67 p=0.035). **G-I.** Heatmap of SERT (**top**) and Gad2 (**bottom**) activity time-locked Trial Start (G), Decision Making (H), and Reward Acquisition (I) for one representative mouse across no-effort sessions. Each row represents an individual trial, with variable trial durations. Black areas indicate NaN values corresponding to periods after trial termination. Line plots represent mean (line) ±SEM (shaded area). Paired line plots represent individual datapoints connected by lines between pre and post.

We next tested whether these phasic responses to each task variable are modulated by reward choice and effort requirements when choosing between effortful high-reward and effortless low-reward options. To address these questions, we compared phasic responses in DRN^SERT^ and DRN^Gad2^ neurons on high-versus low-reward trials, stratified by effort requirements, during the decision making and reward acquisition phases of the task. In the first block of sessions, when no effort was required and choices were driven by simple reward discrimination, DRN^SERT^ and DRN^Gad2^neurons showed similar responses during high- and low-reward trials, both at decision making (**Figure 4A-B**) and reward acquisition (**Figure 4C-D**). This indicates that SERT and GABA signals are not sensitive to the relative reward value alone.

**Figure 4:**
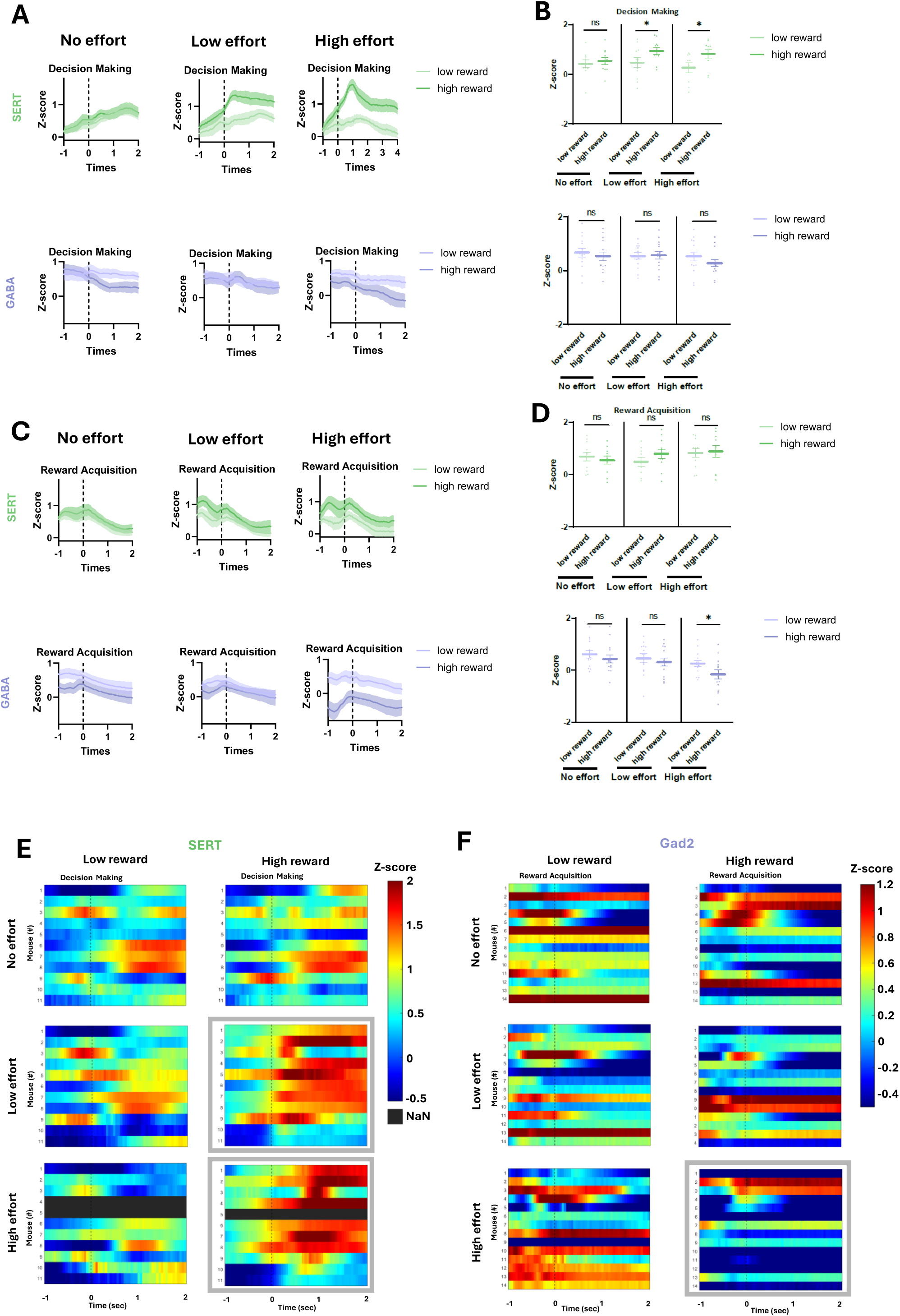
Differential phasic activity associated with reward choice across effort levels. **A.** Averaged traces of SERT (**top**) and Gad2 (**bottom**) neurons around Decision Making for high- and low-reward trials across no-effort, low-effort and high-effort sessions. **B.** Quantification of the signal difference between high- and low-reward trials during Decision Making. SERT activity in higher during high-reward trials in comparison to low-reward trials for low effort and high effort sessions, but not for no effort sessions. No difference of activity is found in Gad2 neurons between high- and low-reward trials during Decision Making regardless of the level of effort (n=11 SERT mice, n=14 Gad2 mice, Mixed-effects analysis: Effort effect p=0.463, Reward effect p=0.109, Neuronal type effect p=0.705, Reward*Neuronal type effect p=0.002, Effort*Reward*Neuronal type effect p=0.113, Bonferroni corrected post-hoc for high versus low reward trials: SERT no effort p>0.999, SERT low effort p=0.023, SERT high effort p=0.020, Gad2 no effort p>0.999, Gad2 low effort p>0.999, Gad2 high effort p=0.388). **C.** Averaged traces of SERT (**top**) and Gad2 (**bottom**) neurons around Reward Acquisition for high- and low-reward trials across no-effort, low-effort and high-effort sessions. **D.** Quantification of the signal difference between high- and low-reward trials during Reward Acquisition. No difference of activity is found in SERT neurons between high- and low-reward acquisition, regardless of effort requirement. Gad2 activity in lower during high-reward trials in comparison to low-reward trials for high-effort sessions, but not for no-effort or low-effort sessions (n=11 SERT mice, n=14 Gad2 mice, Mixed-effects analysis: Effort effect p=0.120, Reward effect p=0.596, Neuronal type effect p=0.015, Reward*Neuronal type effect p=0.003, Effort*Reward*Neuronal type effect p=0.189, Bonferroni corrected post-hoc for high versus low reward trials: SERT no effort p>0.999, SERT low effort p>0.999, SERT high effort p=0.529, Gad2 no effort p=0.676, Gad2 low effort p>0.999, Gad2 high effort p=0.011). **E.** Heatmap of z-scored SERT activity per mouse time-locked to Decision Making. Each row represents an individual mouse. Black areas indicate NaN values corresponding to reward–effort conditions that were not experienced by the mouse during the session. Gray boxes highlight effort conditions showing increased signal on high-reward trials compared to low-reward. **F.** Heatmap of z-scored Gad2 activity per mouse time-locked to Reward Acquisition. Each row represents an individual mouse. Gray box highlights effort condition showing decreased signal on high-reward trials compared to low-reward. Line plots represent mean (line) ±SEM (shaded area). Swarm plots represent mean (line) ±SEM and individual datapoints (dots).

Instead, a signal discriminating between high- and low-reward choices emerged only when reward evaluation occurred in the presence of differential effort requirements (**Figure 4C-D**)., In the later sessions when an effort component (low then high effort) was added to the decision-making process, activity in DRN^SERT^ and DRN^Gad2^ neurons show a differential activity between high-reward and low-reward choices that was specific to certain epoch of the task. The SERT signal, time-locked to decision making, was higher when mice chose the high-reward option in comparison to the low reward during effortful sessions, regardless of the magnitude of the effort required (**Figure 4A-B**). These differences were not observed during reward acquisition (**Figure 4C-D**). In contrast, the GABA signal time-locked to decision making did not differ between high-reward and low-reward choices regardless of effort requirements (**Figure 4A-B**), whereas the GABA signal, time-locked to reward acquisition, was lower when mice were consuming the high reward, but in the high-effort condition (**Figure 4C-D**). These differential signals between high- and low-reward trials were consistent across mice (**Figure 4E-F**). These findings indicate that the decision-related phasic response of DRN^SERT^ neurons scales with effort. Conversely, the reward acquisition-related decrease in DRN^Gad2^ activity was more pronounced following high-effort choices vs. low-effort choices. Together, these findings show that effort selectively enhances decision- and reward acquisition-related signals in DRN neurons in a cell type-specific manner, suggesting that these neuronal population encode reward value as function of effort costs rather than objective reward value alone.

Together, these findings reveal that DRN^SERT^ and DRN^Gad2^ neurons differ in both their tonic and phasic coding of effort-based decisions. We next investigated the relative contribution of these two activity components to behavioral choice in the effortful conditions by training an SVM decoders using tonic activity alone, phasic activity alone, or both signals combined (**Figure 5A**). The Tonic decoder used the mean neuronal activity across the entire trial as a single predictor. The Phasic decoder used a single predictor, defined as the mean neuronal activity during the task epoch in which low- and high-reward signals diverged, corresponding to the decision-making period for DRN^SERT^ neurons and the reward-acquisition period for DRN^Gad2^ neurons. The Full decoder included both measures as independent predictors, allowing us to assess how tonic and phasic activity jointly contributed to behavioral choice (see Methods for details). In DRN^SERT^ neurons, we found that a full model combining tonic and phasic responses of high-effort trials could predict whether mice selected the high- or low-reward option (**Figure 5B**), with significantly greater accuracy than a model trained on shuffled data. Interestingly, the models including only tonic or phasic signals failed to predict reward choices at rates greater than chance, suggesting that tonic and phasic signals of DRN^SERT^ neurons carry distinct information for predicting effort-based decision-making. In contrast in DRN^Gad2^ neurons, although the full model had a higher accuracy, the model containing only phasic response could successfully predict behavioral choice at rates greater than chance (**Figure 5C**). This result brings new evidence regarding the limited influence of tonic variations in GABA activity on the coding of effortful reward seeking and the specificity of the cell type on the coding of effort-based decision making.

**Figure 5:**
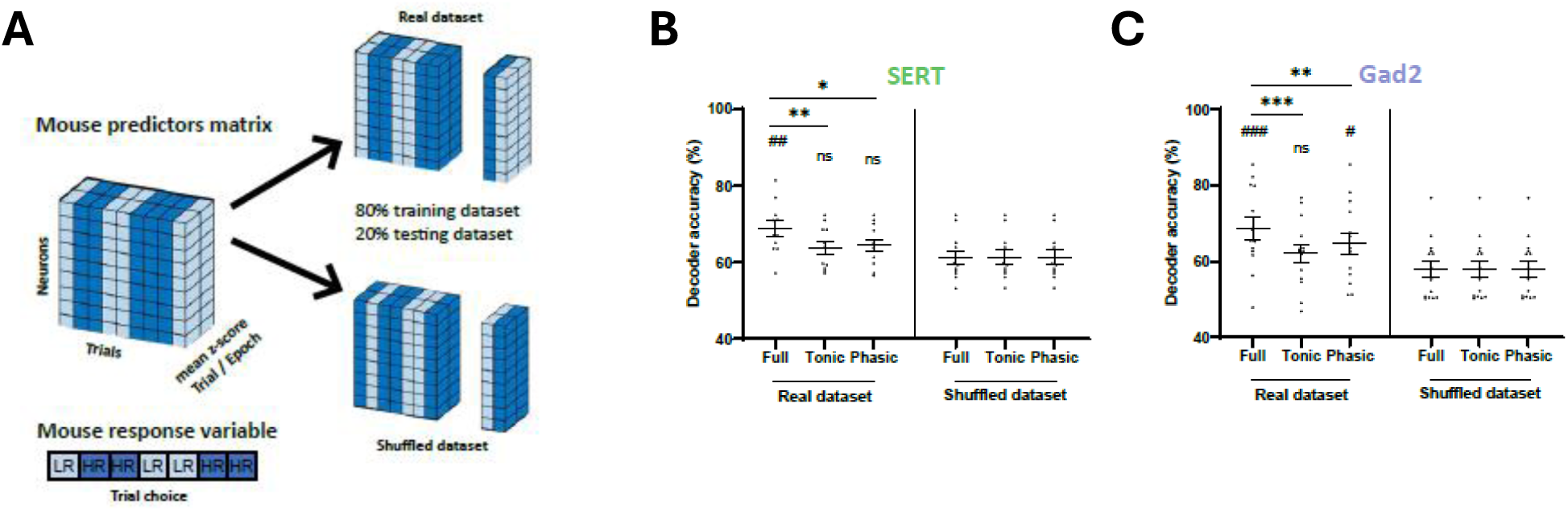
Decoding of reward choices based on phasic and tonic activity of DRN^SERT^ and DRN^Gad2^ neurons. **A.** Schematic of the Full (Tonic + Phasic), Tonic-only, and Phasic-only SVM models and the corresponding shuffled control dataset. **B.** Decoder accuracy of the different SVM models for predicting reward choice from DRN^SERT^ activity using the real and shuffled datasets. Only the Full (tonic + phasic) model successfully decoded reward choice above chance level. (n=11 mice, Mixed-effects analysis: Model effect p=0.014, Dataset effect p<0.001, Model*Dataset effect p=0.017, # Bonferroni corrected post-hoc for real dataset versus shuffled dataset : Full p=0.001, Tonic p=0.554, Phasic p=0.306, * Bonferroni corrected post-hoc for model comparisons : Full vs Tonic p=0.005, Full vs Phasic p=0.014). **C.** Decoder accuracy of the different SVM models for predicting reward choice from DRN^Gad2^ activity using the real and shuffled datasets. The SVM decoder successfully predicted reward choice above chance level using the Full (tonic + phasic) and Phasic models. Decoding accuracy was significantly higher with the Full model than with the Phasic model (n=14 mice, Mixed-effects analysis: Model effect p=0.012, Dataset effect p=0.015, Model*Dataset effect p=0.001, # Bonferroni corrected post-hoc for real dataset versus shuffled dataset : Full p<0.001, Tonic p=0.162, Phasic p=0.021, * Bonferroni corrected post-hoc for model comparisons : Full vs Tonic p<0.001, Full vs Phasic p=0.018). Swarm plots represent mean (line) ±SEM and individual datapoints (dots).

Together, these findings DRN^SERT^ and DRN^Gad2^ neurons carry complementary signals during effort-based decision making. DRN^SERT^ neurons carry a decision-related phasic signal whose magnitude increases as a function of both reward magnitude and effort expenditures. In contrast, activity in DRN^Gad2^ neurons is insensitive to the decision-making period in this task but is suppressed following reward acquisition—an effect that is magnified during high-effort sessions. Furthermore, tonic fluctuations in activity in DRN^SERT^ neurons, but not DRN^Gad2^ neurons, modulates behavioral choice persistence on slower time scales. These results suggest that serotonergic and GABAergic DRN populations play distinct but complementary roles in integrating effort and reward information during effortful reward seeking.

## Discussion

Effort-based decision-making requires integrating the benefits of an action with the cost required to obtain it. Although the DRN has been implicated in reward processing, cost-benefit valuation, and physical effort, its contribution to effort-based choice has remained poorly understood. Here, using fiber photometry recordings of genetically defined DRN neuronal populations during an effort-based decision-making task, the present study identifies the DRN as a candidate node for integrating effort cost and reward benefit, complementing previous work implicating prefrontal, striatal, hippocampal, and amygdalar circuits in effort-based choice^26–32^. It reveals cell-type-specific computations where DRN^SERT^ and DRN^Gad2^ populations do not redundantly encode reward choice, but instead carry partially dissociable representations of persistent choice state, action selection, and reward outcome under effort constraints.

A notable finding of this study is that tonic fluctuations in DRN serotonergic activity were selectively associated with persistent high-reward choice behavior. DRN^SERT^ activity was elevated during clustered sequences of high-reward choices relative to clustered low-reward choices, scaled with the length of high-reward choice clusters, and correlated across sessions with the degree of high-reward choice clustering. Together, these observations suggest that serotonergic activity is not simply a trial-by-trial encoding of individual choice value, but instead reflects a broader behavioral state that promotes sustained selection of the high-reward option despite its associated action costs. These resultsare in line with recent proposals that DRN serotonin neurons carry slow signals related to motivational state that support persistent goal-directed behavior over extended timescales, including active reward seeking, waiting and active coping^34,49,50^. Our findings extend this framework to effort-based decision-making. In contrast, DRN GABAergic neurons did not exhibit a comparable tonic relationship with choice persistence, suggesting that their activity is more consistent with transient, phasic responses linked to specific task events dissociated from slow state-dependent representations. Decoding analyses further refined these distinctions between cell types. In DRN^SERT^ neurons, choice could be predicted only when both tonic and phasic activity features were combined, whereas tonic or phasic features alone were insufficient. This suggests that serotonergic choice-related information is embedded within a multiscale signal integrating slow motivational state and faster event-related dynamics, both of which providing behaviorally relevant information. In contrast, in DRN^Gad2^ neurons, phasic activity around reward acquisition carried predictive information about choice, while tonic activity did not improve decoding performance. Although decoding performance does not establish causal necessity, this pattern is consistent with the notion that DRN^SERT^ neurons encode a multiscale signal combining slow motivational state and rapid decision-related components, whereas DRN^Gad2^ activity primarily track discrete task events.

At the phasic level, DRN serotonergic and GABAergic neurons encoded different components of task performance. DRN^SERT^ neurons exhibited a robust signal around the decision epoch that discriminated high-versus low-reward choices during effortful sessions, whereas DRN^Gad2^ neurons did not show comparable choice-selective activity at that time. Instead, DRN^Gad2^ neurons exhibited a differential response at reward acquisition, with a larger decrease in activity for high-reward choices selectively in the high-effort condition. This dissociation suggests that serotonergic and GABAergic DRN populations may participate in different stages of effortful reward seeking. One possibility is that serotonergic neurons preferentially contribute to processes occurring before or during action selection, when the animal commits to overcoming a cost in pursuit of a larger reward, whereas GABAergic neurons may preferentially signal outcome-related information once the costly action has been completed. Although fiber photometry does not allow causal inference or disentangle whether these signals reflect valuation, action commitment or consummatory processing, the temporal dissociation between serotonergic and GABAergic responses points to functional specialization across DRN cell types during effortful reward seeking.

While reward-related signal has been documented in DRN neurons^33–35^, an important feature of the activity patterns recorded here is that neither serotonergic nor GABAergic signals simply tracked reward magnitude in the absence of cost. These functions may reside in other DRN neuronal subtypes or in projection-specific circuits^51,52,16^, and future studies will be needed to elucidate this. During no-effort sessions, when mice discriminated between reward amounts without having to overcome a barrier, both DRN^SERT^ and DRN^Gad2^ responses were similar for high- and low-reward choices. Choice-related difference in neuronal activity emerged only once reward evaluation took place in the presence of effort requirements. This observation argues against the hypothesis that the recorded DRN signals primarily represent reward size per se. Instead, our data are more consistent with the idea that DRN neurons encode reward -related variables in a cost-sensitive manner, when the behavioral significance of a reward depends on the effort required to obtain it. In addition, the signal difference between high- and low-reward options in DRN^SERT^ scales with effort magnitude similarly to cortical neurons involved in computational valuation in effort-based decision-making^29^, suggesting that DRN serotonergic neurons may engage in computational processes similar to those in this task. This computational role of DRN aligns with evidence that serotonergic signaling contributes to multiple dimensions of cost–benefit evaluation, including probability, risk and delayed reward evaluation^38–41^. A theorical model has proposed that DRN serotonergic neurons encode a “beneficialness” signal, reflecting the integration of reward magnitude, probability, and cost (p × R × C)^53^. Our results are consistent with the core idea of this framework in the context of effort-based decision-making, showing that DRN serotonergic activity carries reward-related signals that are shaped by effort cost. However, our data do not directly address the contribution of probabilistic factors to this signal, which will require future experimental investigation.

In summary, this study identifies distinct tonic and phasic signatures of effort-based decision-making in DRN serotonergic and GABAergic neurons. Our results support a model in which DRN^SERT^ activity provides a computational signal linking persistent motivational state to effortful choice, whereas DRN^Gad2^ activity preferentially signals reward acquisition after costly action and may participate in the evaluation or updating of reward outcomes following effort expenditure. Given that local DRN GABAergic neurons provide inhibitory control over serotonergic neurons, thereby shaping DRN output and serotonin release^12^, one possibility is that the reduced reward-related activity of DRN^Gad2^ neurons following costly actions transiently relieves local inhibition of serotonergic neurons, allowing information about reward outcomes after effort expenditure to influence the tonic serotonergic activity associated with persistent selection of the larger reward. However, the DRN also contains a well-characterized population of long-range projecting GABAergic neurons that innervate extra-raphe targets and directly regulate downstream circuits^14^. In the anterior cingulate cortex, selective recordings from local GABAergic interneurons revealed only weak modulation during effort-based decision-making, despite their prominent role in regulating excitatory projection neurons that carry choice-related signals^29^. Together, these observations indicate that the reward-related DRN^Gad2^ signal identified here could reflect either local regulation of serotonergic neurons or activity of long-range projecting GABAergic neurons acting on downstream structures. Future studies dissecting local versus long-range GABAergic subpopulations and determining how they interact with serotonergic ensembles will be essential for understanding how DRN microcircuits and DRN output pathways cooperate to regulate effort-based decision-making.

By elucidating the DRN circuit mechanisms that support effort-based decision-making, the present findings may also inform our understanding of neuropsychiatric and neurological disorders associated with motivational dysfunction, in which the willingness to expend effort for reward is often diminished. In this context, disrupted serotonergic signaling could contribute to these symptoms by weakening the sustained motivational states that support persistent engagement toward larger but more costly rewards, whereas altered DRN GABAergic signaling could impair the evaluation or updating of reward outcomes following effort expenditure. Together, these findings suggest that cell-type-specific dysfunction within the DRN could represent a potential circuit-level mechanism through which effort allocation and reward-seeking behavior become disrupted in psychiatric and neurological disease.

## Materials and Methods

### Animals

Male and female mice were aged 9-12 weeks at the beginning of the experiments. Heterozygous SERT-Flp (B6.Cg-Slc6a4tm1.1(flpo)Luo/J) and Gad2-Cre (B6J.Cg-Gad2tm2(cre)Zjh/MwarJ) mice were used for fiber photometry recoding of DRN serotoninergic and GABAergic neurons respectively. Mice were housed in a Weill Cornell Medical College facility and were maintained on a 12-hour light-dark cycle. Except when water-restricted for the purpose of behavioral training and testing, all mice were given ad libitum access to food and water. Littermates underwent surgeries within the same week, and mice were group-housed with littermates with the same surgery status. All procedures were approved by the Weill Cornell Medicine IACUC.

### Surgery

Animals were placed inside a flow box and anesthetized with isoflurane gas (3% in 1 l/min oxygen) until sedated, at which point they were placed in a stereotax and maintained on 1% isoflurane for the duration of the surgery. Eye gel (Optixcare) was applied to both eyes and body temperature was maintained at 37 °C with a heating pad and rectal probe. Meloxicam (2mg/kg) and buprenorphine (0.5mg/kg) were injected intraperitoneally. Scalp hair was trimmed away, a midline incision was made using fine surgical scissors (Fine Science Tools) for exposing the skull and 0.1ml of local anesthetic agent (bupivacaine 0.25%) was topically applied. Skull was then cleaned with 3% hydrogen peroxide and immediately rinse with saline solution.

1 µl of AAV8-Ef1a-fDIO-GCaMP6s (Addgene #105714) or AAV1.Syn.Flex.GCaMP6s.WPRE.SV40 (Addgene #100845) were injected in the DRN (with respect to the bregma: AP, -4.5 mm; ML, 0; DV, -3.5 mm) of SERT-flp and Gad2-Cre mice respectively. A 4mm optical fiber (Doric Lenses #B280-4401-4) was then implanted in the DRN (with respect to the bregma: AP, -4.5 mm; ML, 1.2; DV, - 3.1 mm; α, +20°) (**Supplementary figure 1A**). Fiber implant was secured with Vetbond (3M) and Metabond.

After surgery, mice received a subcutaneous injection of sterile saline solution and were placed in a recovery cage on a heating pad until they awoke from anesthesia. To avoid post-operative pain, meloxicam was intraperitoneally injected for 3 days.

### Apparatus for effort-based decision-making task

The effort-based decision-making apparatus consists in an elevated T-maze with two reward arms and a central arm. At the entrance of the central arm, IR sensors are used for detecting mice and initiate new trials. At the entrance of the reward arm, custom effort barriers (low-effort barrier – 5cm height, high-effort barrier 10cm height) are positioned to force the animal to climb over them to reach the end of the arm. Feeding needles are mounted through the end walls of the reward arms to allow water delivery. IR sensors are utilized for detecting licking of the reward ports. Animal behavior is video-recorded, and video-tracked using custom scripts.

### Behavioral training

As previously described (Fetcho et al. 2023), the effort-based decision-making task consists in shaping sessions and 3 training phases. Briefly, during shaping sessions, water restricted mice were habituated to drink water through the reward port and to go to the entrance of the central arm to initiate new trials (indicated by a 5 kHz tone). After successfully initiating more than 20 trials within one session (maximum of 20 minutes per session), mice moved to training sessions in which a high reward (high volume of water) was allocated to one reward arm and a low reward (low volume of water) was allocated to the other one. The high reward was assigned to the least preferred arm during the shaping sessions. Mice underwent 3 training phases in which no effort, low effort or high effort (no barrier, low-effort barrier, high-effort barrier) was successively paired with the high reward arm. No effort was ever needed to obtain the low reward. Initially, the ratio of high-reward volume to low-reward volume was 2:1. Animals underwent daily training sessions, with individualized adjustment of the reward ratio until they preferred the high-reward arm in 70% or more of trials within a session for two sessions in a row. Then mice moved to the next training phase with an increase of the effort requirements.

### Fiber photometry recording

Animals with fiber optic implants were attached via patchcord to the photometry system (Neurphotometrics FP3002) prior to training sessions throughout the course of behavioral training. Recordings were acquired for the entirety of each session and were time-locked to the behavioral system thanks to TTL pulses to permit analysis of DRN^SERT^ and DRN^gad2^ activities in relation to effort-based decision-making epochs. Previous studies have reported a notable relationship between DRN neural activity and mice motricity (Kawashima et al. 2016, Nectow et al. 2017). Here, we found a correlation between mice velocity and recorded activity in both the SERT and Gad2 populations, supporting this link (**Supplementary figure 1C**). To avoid misinterpretation of neural activity in the effort-based decision-making task, we regressed the signal recorded by the velocity and acceleration of each individual mice. Velocity and acceleration used for the regression were measured during the intertrial interval, as the mice returned to the central arm to initiate the next trial.

### Fixed tissue processing and imaging

To verify the location of the optical fiber, animals were transcardially perfused with ice cold 1x PBS and 4% paraformaldehyde (PFA). The brains were dissected, post-fixed overnight in 4% PFA at 4°C, cryoprotected in 20% sucrose, frozen rapidly, and then stored at -20°C before coronal sectioning with a cryostat (40 mm thick). Samples were mounted on 25x75x1 mm slides (Fisher Superfrost) with DAPI-infused Fluoroshield mounting medium.

Images were carrying out on the Leica DM 5500 B microscope with a Leica DFC360 FX fluorescent camera (7x magnification) and EL600 light source, using a 10x apochromat objective with an NA of 0.4. Images were taken in two channels - a green channel to capture GFP and a blue channel to capture DAPI.

### Binary decoding with SVM classifiers

MATLAB’s *fitcsvm* function was used to test the ability to decode, from different measures of SERT and Gad2 fiber photometry recording, the mice choice between high reward and low reward side. The models were trained using SVM decoder with 5-fold cross-validation. On each iteration, data were separated (bootstrapped) into training (80% of trials) and testing (20% of trials) sets and in each set, trials were separated into equal numbers from each binary value for the feature of interest. Decoding accuracy was expressed as percentage of correct choice prediction per mouse. We compared the accuracy of the decoder depending on whether we used as predictors the tonic coding (mean trial activity), the phasic coding (mean activity around decision making or reward acquisition for SERT or Gad2 neurons respectively) or a combination of both. To test the ability of the model to randomly decode the mice choice, we shuffled (1 000 iterations) the choice of the training dataset using *randswap* and tested the model on a test dataset with real values.

### Statistics

All statistical analyses were performed using Matlab and Prism (GraphPad). For comparison between 2 datasets paired Wilcoxon tests were used. For comparison between more than 2 datasets, Mixed Model analysis was performed with mouse identity as random effect Bonferroni corrections were performed for multiple post hoc comparisons. Correlations were performed using Spearman test. Significant result was set at p < 0.05. No statistical methods were used to predetermine sample sizes, but our sample sizes were similar to those reported in previous publications^29^.

---

**Supplementary figure 1:**
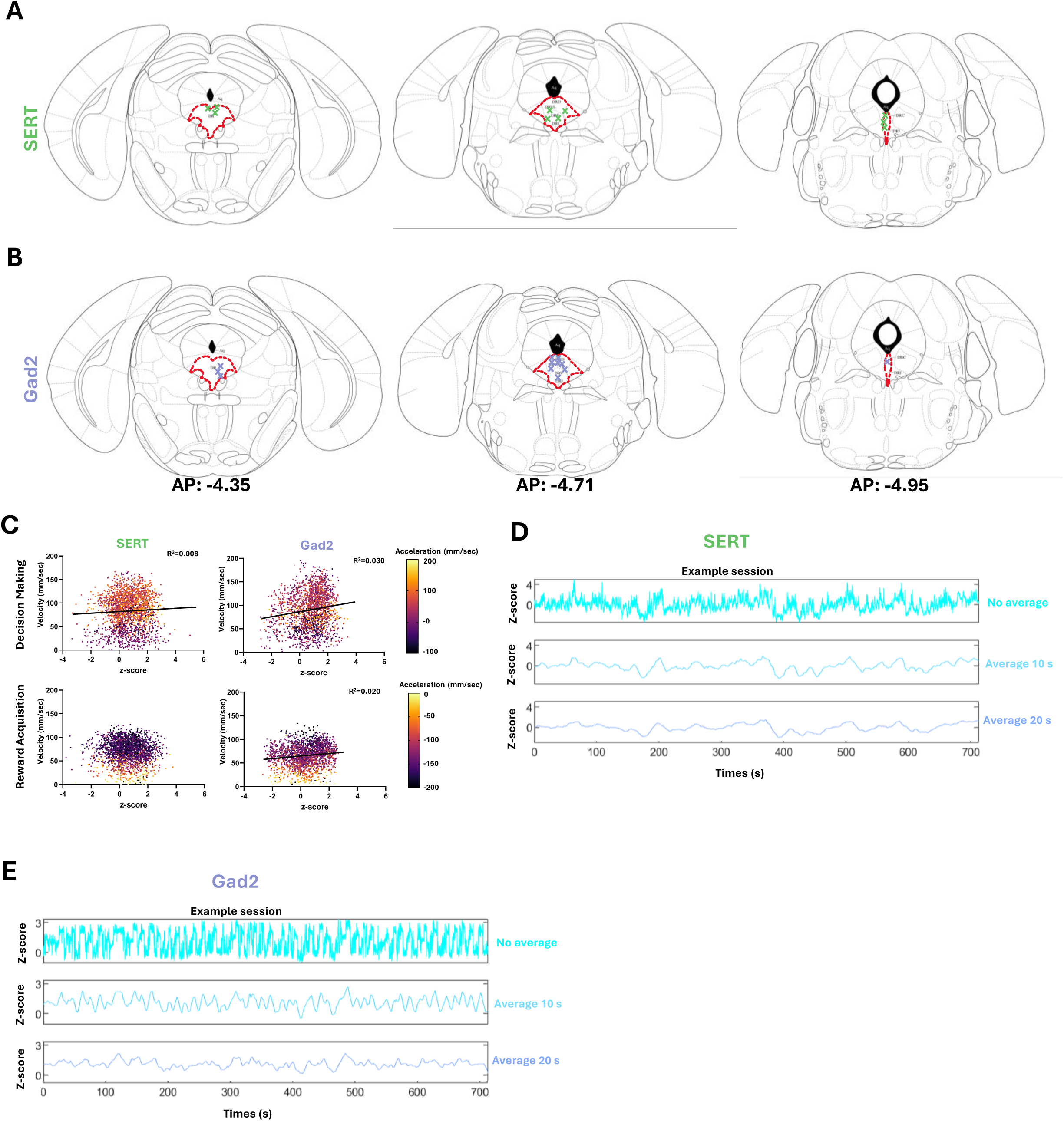
**A.** Optical fiber placement for fiber photometry recording of SERT neuronal population. **B.** Optical fiber placement for fiber photometry recording of Gad2 neurons. **C.** Correlation between neuronal population activity and mice motricity. SERT neuronal activity is positively correlated to mice velocity during decision making but not during reward acquisition (**left**, n=2150 no effort trials, Multiple Linear Regression: Decision making Velocity effect p=0.035 Acceleration effect p=0.468 Velocity*Acceleration effect p=0.440, Reward acquisition Velocity effect p=0.215 Acceleration effect p=0.537 Velocity*Acceleration effect p=0.976). Gad2 neuronal activity is positively correlated to mice velocity during decision making and reward acquisition (**right**, n=1891 no effort trials, Multiple Linear Regression: Decision making Velocity effect p<0.001 Acceleration effect p=0.451 Velocity*Acceleration effect p=0.995, Reward acquisition Velocity effect p=0.003 Acceleration effect p=0.782 Velocity*Acceleration effect p=0.836). **D.** Example session recording of a SERT neuronal population. Moving averages of neuronal activity over 10- and 20-second windows, highlighting slow fluctuations in population activity. **E.** Example session recording of a Gad2 neuronal population. Moving averages of neuronal activity over 10- and 20-second windows, highlighting slow fluctuations in population activity. Scatter plots represent individual datapoints (dots).

